# RxRx3: Phenomics Map of Biology

**DOI:** 10.1101/2023.02.07.527350

**Authors:** Marta M. Fay, Oren Kraus, Mason Victors, Lakshmanan Arumugam, Kamal Vuggumudi, John Urbanik, Kyle Hansen, Safiye Celik, Nico Cernek, Ganesh Jagannathan, Jordan Christensen, Berton A. Earnshaw, Imran S. Haque, Ben Mabey

## Abstract

The combination of modern genetic perturbation techniques with high content screening has enabled genome-scale cell microscopy experiments that can be leveraged to construct *maps of biology*. These are built by processing microscopy images to produce readouts in unified and relatable representation space to capture known biological relationships and discover new ones. To further enable the scientific community to develop methods and insights from map-scale data, here we release **RxRx3**, the first ever public high-content screening dataset combining genome-scale CRISPR knockouts with multiple-concentration screening of small molecules (a set of FDA approved and commercially available bioactive compounds). The dataset contains 6-channel fluorescent microscopy images and associated deep learning embeddings from over 2.2 million wells that span 17,063 CRISPR knockouts and 1,674 compounds at 8 doses each. **RxRx3** is one of the largest collections of cellular screening data, and as far as we know, the largest generated consistently via a common experimental protocol within a single laboratory. Our goal in releasing **RxRx3** is to demonstrate the benefits of generating consistent data, enable the development of the machine learning methods on this scale of data and to foster research, methods development, and collaboration.

For more information about **RxRx3** please visit RxRx.ai/rxrx3

## Introduction

Advances in genomic technologies and high content screening (HCS) capabilities have enabled unbiased, large-scale expression and image-based profiling experiments of genetic perturbations. These screens have massive potential to uncover novel biology and accelerate drug discovery, however known issues prevent these screens from obtaining this potential. For example, most publications to date use pooled CRISPR screens which read out counts of CRISPR guide RNAs (gRNAs) introduced into cells in bulk prior to surviving a biological challenge, yet these selection-based readouts are limited in that they cannot differentiate between genes acting via different mechanisms. To overcome this limitation, high content transcriptome profiles have been measured from pooled CRISPR screens using single-cell RNA sequencing (scRNA-seq)^1^. While genome-wide scRNA-seq experiments are feasible^2^, scRNA-seq is still prohibitively expensive at a genome-wide scale and experiments are typically limited to a few thousand target genes^1^. Additionally, their pooled nature makes it impossible to control the number of cells transfected with each gRNA and to account for cell-cell interactions arising from different pairs of perturbations.

Arrayed genetic screening is an alternative to pooled screening that overcomes these limitations. In arrayed genetic screens, populations of cells are perturbed by a single gRNA in isolation, for each gRNA in the collection^1^. For example, the JUMP-CP Consortium recently released a cellular imaging dataset of cells under ~116,000 chemical and ~15,000 genetic perturbations^3^. Due to the size of this dataset (~800,000 images), the consortium found it necessary to produce these images across 12 different data-generating centers, which could result in the confounding of these readouts by cross-laboratory batch effects.

In order to create the highest degree of consistency between the readouts of such large datasets, Recursion (recursion.com) has scaled the use of arrayed CRISPR-mediated gene knockouts to produce genome-wide perturbation datasets using cellular morphological imaging as readouts^4^ within a single automated laboratory. As an example of such readouts, in this paper we describe **RxRx3**, a publicly available genome-wide arrayed CRISPR knockout screen, containing over 2.2 million 6-channel fluorescent images (using a modified Cell Painting protocol^5,6^) of 17,063 CRISPR knockouts and 1,674 compounds at multiple doses (Figure 1). Our intention in releasing RxRx3 is to demonstrate the benefits of generating consistent data in this way, as well as enable researchers to develop the next generation of machine learning methods that leverage these datasets in building *maps of biology^7^* to power the discovery of novel relationships between genes and compounds.

**Figure 1:**
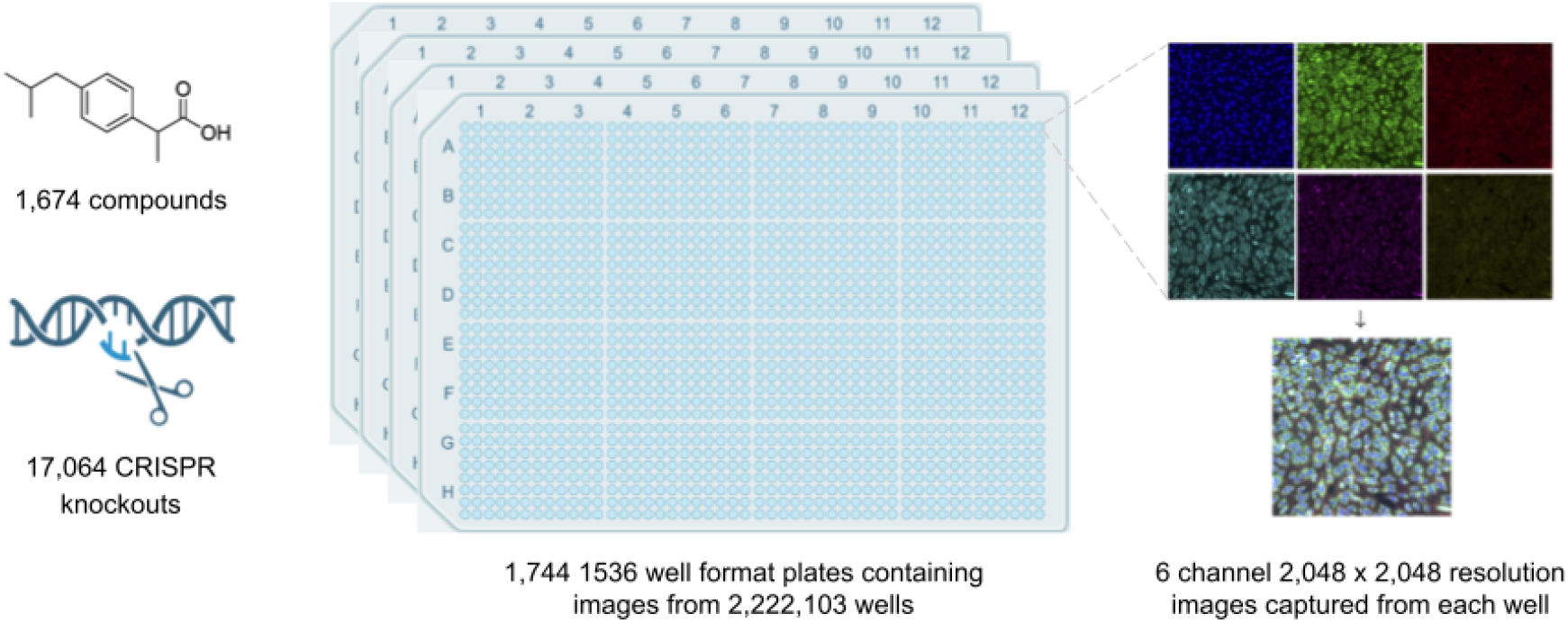
RxRx3 overview. HUVEC cells are plated in 1,536-well plates and treated with one of 17,063 CRISPR knockouts or 1,674 compounds. Six individual channels are captured via staining with Hoechst 33342 (channel 1, blue), Alexa Fluor 488 Concanavalin A (channel 2, green), Alexa Fluor 568 Phalloidin (channel 3, red), Syto14 (channel 4, cyan), MitoTracker Deep Red FM (channel 5, magenta), Alexa Fluor 555 Agglutinin (channel 6, yellow). The similarity in content between some channels is due in part to the spectral overlap between the fluorescent stains used in those channels.

## Biological discovery use cases

**RxRx3** was designed to discover novel gene-gene, gene-compound, and compound-compound relationships. As every experiment in the set was executed under nearly identical conditions (aside from nuisance variations related to experimental batch effects and differences between compound and CRISPR protocols), images of each perturbation can be directly compared with images from any other perturbation in the dataset. The primary computational challenge when working with **RxRx3** is featurizing the images in a way that is invariant to nuisance batch variations (which can be significant in the raw imaging data) but preserves the biologically meaningful information for comparing effects of perturbations. Typically cell segmentation followed by feature extraction is used to quantify every cell in the experiment and then QC and batch correction techniques are applied^8^. More recently, deep learning-based embedding models have been shown to be a powerful alternative to segmentation and feature extraction pipelines for featurizing microscopy images^9,10^. To make **RxRx3** useful to a variety of users, we released the raw images as well as pre-computed 128-dimensional embeddings (from a proprietary deep learning model) for every well in the experiments.

With features computed for **RxRx3**, whether from the included embeddings or another high-quality featurization method, one can use them to identify relationships between biological entities (e.g., gene-gene interactions arising from protein-protein interactions, protein complexes or signaling pathways)^7^. RxRx3 also explores the interface between biology and chemistry where the 1674 compounds can be used to search for relationships between genes and compounds across 8 doses. These can be inferred by computing distances (e.g., Euclidean or cosine) between aggregated perturbation representations, where smaller distances implies a greater degree of similarity between the underlying cellular conditions. These distances can in turn be used to visualize the global structure of perturbations through further dimensionality reduction techniques such as uniform manifold approximation (UMAP) or direct comparison, such as we demonstrate with **MolRec™^11^**. More details about these use cases can be found in Celik et al., 2022^7^.

## Machine learning use cases

**RxRx3** will enable the machine learning and HCS communities to develop new methods for analyzing such datasets. Deep learning approaches have come to dominate nearly all computer vision tasks^12,13^. Despite this trend, the vast majority of high-content screening studies use traditional segmentation and feature extraction analysis pipelines^14^. Adoption of deep learning in the HCS community has been hindered by the fact that different labs have very different experimental protocols, including differences in cell types, imaging hardware, staining protocols, reagents, etc. As a result, it can be counterproductive or even impossible to reuse a model pre-trained on one dataset with a different dataset. One notable exception is cell and nuclear segmentation, for which similar imaging protocols are used across labs, and deep learning models have been developed that can generalize to new datasets without additional tuning^15–17^. Despite this development, segmented cells still need to be quantified with existing feature extraction approaches.

Attempts to quantify cellular phenotypes with deep learning have been limited by the lack of datasets with high-quality labels. For some specific tasks like protein localization^18,19^ or cell cycle classification^20^, enough cellular data has been manually annotated to train high-quality supervised deep learning models on specific datasets. However, these applications represent a small set of the possible phenotypes that can be characterized with HCS. We, along with other members of the HCS community, aspire to build unbiased deep learning analysis pipelines that can detect novel biological phenotypes that have not or cannot be manually annotated.

Progress toward general purpose deep learning frameworks for HCS have been driven by weakly-supervised learning and more recently by self-supervised learning (SSL). In weakly-supervised learning^9,21^, perturbation metadata (i.e. the perturbation applied to each well) is used to label HCS images and train supervised deep learning models. This approach is currently the state-of-the-art approach for many use cases. Its strength stems from learning common patterns that exist in cells exposed to the same perturbation but imaged in different experimental batches. Its weakness, however, is in requiring several experimental batches of each perturbation to learn performant models, and from uncertainty over which perturbations should be selected for model training (Nikita, et al.^9^ suggest using classical features to select perturbations with stronger phenotypic effects for training).

A more recent alternative deep learning framework that has been successful at learning HCS phenotypes is self-supervised learning^22,23^. These models work by learning to relate cropped and augmented regions from the same input image, thereby learning the patterns that are inherent in the dataset. In the computer vision community, these types of approaches are now state-of-the-art, but often benefit from pre-training on datasets that are orders of magnitude larger that task-specific datasets like ImageNet. **RxRx3** can easily be used for training weakly-supervised models since the treatment column in the metadata has a unique string for each of the 102,705 perturbations in the dataset. Additionally, **RxRx3** can be used for training SSL models by leveraging the raw images directly and potentially using relevant metadata^23–25^.

## Methods

### Dataset description

**RxRx3** contains 6-channel fluorescent microscopy images of HUVEC cells stained with a modified cell painting protocol^5,6^ from 2,222,103 experimental wells. Perturbations in these wells span 17,063 CRISPR knockouts and 1,674 commercially available bioactive compounds at 8 concentration. Most CRISPR knockouts have 6 unique guide RNAs targeting the gene. The identities of 736 genes are unblinded to allow discovery and validation work, whereas the identities of the remainder are anonymized. The dataset statistics are summarized in the table 1 below:

**Table 1.**

|  |  |
| --- | --- |
| Images (individual fluorescent channel *png* files) | 13,332,618 |
| Wells | 2,222,103 |
| Plates (1536 format) | 1,744 |
| Experiments (each containing multiple plates) | 180 |
| Number of unique compounds | 1,674 |
| Number of unique CRISPR gene targets | 17,063 |
| Identified unique CRISPR gene targets | 736 |
| Anonymized unique CRISPR gene targets | 16,328 |
| Number of unique CRISPR guide RNAs | 101,029 |

**CRISPR knockout experiments (gene001 - gene176**) contain 9 plates and each plate contains control and target guides. Wells that have not passed quality control specifications have been left out of the dataset.

**Compound experiments (compound001 - compound004)** contain 1,674 compounds at 8 concentrations and are typically repeated 4 times (more often for control compounds). Wells that have not passed quality control specifications have been omitted from the dataset.

### Deep learning embeddings

All images were featurized to 128 dimensions using a custom trained neural network similar to methods previously reported by Recursion^10^.

### Data acquisition

#### Cells

HUVEC (Human umbilical vein endothelial cells, Lonza, C2519A) were cultured according to manufacturer’s recommendations in EGM2 (Lonza, CC-3162). CRISPR-Cas9 reagents were purchased from Integrated DNA Technologies, Inc. (IDT) and prepared following the manufacturer’s guidelines and protocols.

Cells that received compounds were treated with compound via acoustic transfer (Echo 555, Labcyte) at final concentrations of 0.003, 0.01, 0.03, 0.1, 0.3, 1, 3, 10 uM for 18-24 hrs. Cells were seeded into 1536-well microplates (Greiner, 789866) via Multidrop (Thermo Fisher) and incubated at 37C in 5% CO2 for the duration of the experiment. Plates were stained using a modified cell painting protocol^5,6^. Cells were treated with mitotracker deep red (Thermo, M22426) for 35m, fixed in 3-5% paraformaldehyde, permeabilized with 0.25% Triton X100, and stained with Hoechst 33342 (Thermo), Alexa Fluor 568 Phalloidin (Thermo), Alexa Fluor 555 Wheat germ agglutinin (Thermo), Alexa Fluor 488 Concanavalin A (Thermo), and SYTO 14 (Thermo) for 35 minutes at room temperature and then washed and stored in HBSS+0.02% sodium azide.

#### Imaging

Plates were imaged using Image Express Micro Confocal High Content Imaging System (Molecular Devices) microscopes in widefield with 10X objectives. Single sites per well were acquired with 6 channels per well filters as previously specified^6^.

